# Critical role for the TGF-β1/mTORC1 signalling axis in defining the transcriptional identity of *CTHRC1*+ pathologic fibroblasts

**DOI:** 10.1101/2024.10.12.617979

**Authors:** Jo-Anne AM Wilson, Rachel Walters, Delphine Guillotin, Naftali Kaminski, Silvia Parolo, Manuela Platé, Rachel C Chambers

**Affiliations:** Centre for Inflammation and Tissue Repair, UCL Respiratory, Rayne Building, University College London, London, WC1E 6JF, UK; Section of Pulmonary, Critical Care and Sleep Medicine, Yale School of Medicine, New Haven, CT, USA; Fondazione The Microsoft Research - University of Trento Centre for Computational and Systems Biology (COSBI), Rovereto, Italy

## Abstract

Fibrosis, defined as the abnormal deposition of extracellular matrix (ECM), represents the concluding pathological outcome in a number of inflammatory, immune-mediated and metabolic diseases. Recent single cell RNA-sequencing studies have highlighted the diversity and functional heterogeneity of fibroblast populations in multiple fibrotic conditions. These include a novel pathogenic population of high collagen-producing fibroblasts, characterised by expression of the secreted glycoprotein, collagen triple helix repeat containing 1 (CTHRC1). The cardinal pro-fibrotic mediator TGF-β1 has been widely implicated in promoting fibrogenesis and is a potent inducer of *CTHRC1* and *COL1A1* expression. In addition to the canonical Smad signalling pathway, TGF-β1-induced collagen I production is under critical regulatory control by the mTORC1/4E-BP1 signalling hub. Using pharmacological inhibition in combination with gene-editing approaches, we now demonstrate that the role of the mTORC1 axis extends to the regulation of over a third of all TGF-β1 regulated matrisome genes. We provide further evidence that the global transcriptome of TGF-β1-stimulated fibroblasts *in vitro* matches that of a subpopulation of the high collagen expressing *CTHRC1*+ pathological fibroblast population in the IPF lung. In contrast, the TGF-β1 induced transcriptome of fibroblasts in which mTORC1 signalling is disrupted (by *RPTOR* gene editing using CRISPR-Cas9) does not map to any fibroblast population present in human control or fibrotic lung. Using the novel and selective mTORC1 inhibitor RMC-5552, we further demonstrate a direct functional link between mTORC1 signalling and the acquisition of key marker genes which define the *CTHRC1+* fibroblast population in IPF. These data demonstrate, for the first time, a critical role for the mTORC1 signalling hub in determining the transcriptional identity of the *CTHRC1+* pathological fibroblast population and provide strong scientific support for targeting mTORC1 as a therapeutic strategy in IPF and potentially other fibrotic conditions associated with dysregulated TGF-β1 signalling in the fibrotic niche.

## Introduction

Fibrosis is a highly conserved tissue injury response and represents the concluding pathological outcome in several common chronic inflammatory, immune-mediated and metabolic diseases. Pathological fibrosis can affect any organ and accounts for up to 45% of deaths in the industrialised world (1). Despite this, therapeutic options remain limited. In terms of rapid progression to organ failure, idiopathic pulmonary fibrosis (IPF) remains one the most rapidly progressive and fatal fibrotic conditions. In spite of the introduction of the first approved anti-fibrotic agents, pirfenidone and nintedanib, life expectancy remains at 3-5 years from diagnosis as these agents slow but do not halt disease progression (2–4). Fibrotic foci (also known as fibroblastic foci) represent the cardinal pathogenic lesions in IPF and comprise activated fibroblasts, the key effector cells responsible for dysregulated extracellular matrix deposition. While the aetiology of IPF remains elusive, current evidence suggests that this condition arises as a consequence of a combination of genetic risk factors and environmental exposures and manifests in aged individuals as a highly abnormal wound healing response characterised by dysregulated epithelial-mesenchymal crosstalk (5). Recent single cell RNA-sequencing (scRNA-Seq) studies have uncovered the diversity and functional complexity of fibroblast subtypes and have identified multiple fibroblast subpopulations which are only present in fibrosis (6, 7). Notably, this includes a recently identified population of high collagen producing fibroblasts, characterised by high expression of the secreted glycoprotein, collagen triple helix repeat containing 1 (CTHRC1) which plays a pivotal role in ECM deposition and wound repair (8–10).

Although the triggers of the aberrant fibrogenic response vary between tissues, there are core mechanisms shared across multiple organs (11–13) This includes persistent signalling by transforming growth factor β1 (TGF-β1), the most potent fibrogenic cytokine characterised to date. TGF-β1 controls the activation of fibroblasts and promotes extracellular matrix (ECM) deposition by upregulating the expression of an extensive pro-fibrotic transcriptional programme involving multiple matrisome and matrisome associated genes (14). More recently, CTHRC1 has been reported to be a primary target for TGF-β1 signalling in the fibrotic microenvironment in several fibrotic conditions (15).

TGF-β1 signals through the canonical Smad pathway and several non-canonical signalling pathways and recent evidence from our laboratory revealed that the canonical Smad pathway cooperates with mTOR signalling to mediate the potent fibrogenic effects of TGF-β1 on collagen I deposition in primary lung fibroblasts derived from control subjects and IPF patients as well as in cancer associated fibroblasts (lung adenocarcinoma), hepatic stellate cells and skin fibroblasts (16). mTOR is a nodal serine/threonine kinase that is present in two similarly organised but functionally distinct signalling complexes, mTOR complex 1 (mTORC1) and 2 (mTORC2). The two mTOR complexes share several accessory proteins (Deptor, MLST8, Tti1, Tel2) with mTORC1 also containing Raptor and PRAS40 and mTORC2 containing Rictor, Protor1/2 and mSin1. The unique accessory subunits endow each complex with distinct modes of substrate recognition and regulation, which in turns dictates their downstream cellular activities. mTORC1 integrates signals such as growth factors and the availability of nutrients, energy and oxygen, and triggers cellular responses including protein translation, lipid biosynthesis, senescence and autophagy, to either boost anabolism or suppress catabolism via the phosphorylation of ribosomal protein S6 kinase beta-1 (S6K1), also known as p70S6 kinase (p70S6K) and eukaryotic translation initiation factor 4E-binding protein 1 (4E-BP1), amongst others. mTORC2 is responsive to growth factor signalling and its functions include the regulation of cell proliferation, survival and cytoskeletal remodelling via the activation of protein kinase B (AKT), serum and glucocorticoid-induced protein kinase 1 (SGK1) and protein kinase C (PKC).

The mTOR axis has been implicated in multiple cancers, as well as diabetes and neurodegenerative disease and this axis has been avidly pursued as a therapeutic target in these disease settings. First generation allosteric inhibitors, such as rapamycin and its analogues (rapalogs), bind to the mTOR FKBP-rapamycin binding domain (FRB) and prevent the phosphorylation activity of mTORC1 with preferential impact on the phosphorylation of its major downstream substrate, p70S6K (16). Due to its active site being partially blocked by Rictor, mTORC2 is unaffected unless exposure is prolonged (17). Second and third generation mTOR inhibitors comprise dual mTORC1 and mTORC2 ATP-competitive mTOR inhibitors. These function by blocking the mTOR ATP-binding pocket and are currently being explored for their potential in treating various cancers (18). More recently, the generation of a novel class of selective bi-steric inhibitors of mTORC1, which interact with both the orthosteric and the allosteric binding sites in order to deepen the inhibition of mTORC1 while also preserving selectivity for mTORC1 over mTORC2, have been developed. These mTORC1 selective inhibitors potently inhibit p70S6K and 4E-BP1 phosphorylation, while limiting unwanted effects on glucose metabolism and providing relief of AKT-dependent feedback inhibition of receptor tyrosine kinase (RTK) expression as a result of concomitant mTORC2 inhibition (19). At the time of writing, the lead compound, RMC-5552, is a clinical-phase asset in the oncology setting (20). The aim of the current study was to investigate the role of the mTOR signalling hub in regulating the global transcriptional response to TGF-β1 and further whether mTOR contributes to the transcriptional identity of a high collagen expressing *CTHRC1*+ fibroblast population.

## Results

### ATP-competitive inhibition of mTOR attenuates TGF-β1-induced extracellular matrix gene expression

We have recently provided strong support for mTORC1 as a key signalling node in regulating TGF-β1-induced collagen I production (16). Recent scRNA-Seq studies have identified several pathological fibroblast subpopulations which are present in multiple fibrotic conditions, including IPF (6, 7). However, whether mTORC1 acts as a master transcriptional regulator of the ECM and plays a role in shaping the functional complexity of fibroblast subpopulations, including the acquisition of the *CTHRC1+* fibroblast phenotype in response to TGF-β1 signalling in the fibrotic niche, is not known. To gain further insight into these questions, we first examined an existing RNA-Seq dataset from our laboratory (GSE102674) using the DESeq2 package to conduct an interaction analysis comparing the effect of dual mTOR inhibition using the ATP-competitive mTOR inhibitor, AZD8055 with the partial mTORC1 inhibitor, rapamycin, on the TGF-β1-induced transcriptome in primary human lung fibroblasts (pHLFs).

Differential gene expression analysis of pHLFs, untreated or stimulated with TGF-β1, identified 4,911 genes that were significantly modulated in response to TGF-β1, of which, 2,258 genes were significantly (adjusted p-value < 0.05) upregulated with fold change > 1.5 and 2,653 genes were significantly downregulated with fold change < -1.5. These genes are listed in the interactive Supplementary Table 1 to enable readers to run their own future queries on our datasets. Within the TGF-β1 responsive genes, we identified 1,203 genes that exhibited sensitivity to AZD8055 through DESeq2 interaction analysis (Figure 1a) indicating that up to 24.5% of genes impacted by TGF-β1 are likely under mTOR regulatory control (for a full list please see interactive Supplementary Table 2). Pathways analysis of these differentially expressed genes (DEGs) revealed that the most over-represented AZD8055-sensitive pathways were critically linked to the fibrotic response (Figure 1b). The top pathway, ‘Extracellular matrix organisation’, encompassed more than 60 ECM and ECM-related genes. Additionally, pathways associated with Collagen formation and biosynthesis, Elastic fibre formation and ECM interactions ranked among the top 15. The pathway ‘ATF4 activates genes in response to endoplasmic reticulum stress’ was also significantly enriched. Furthermore, critical ECM genes including collagens, proteoglycans and glycoproteins were upregulated by TGF-β1 and counter-modulated by AZD8055 treatment (Figure 1c).

**Figure 1:**
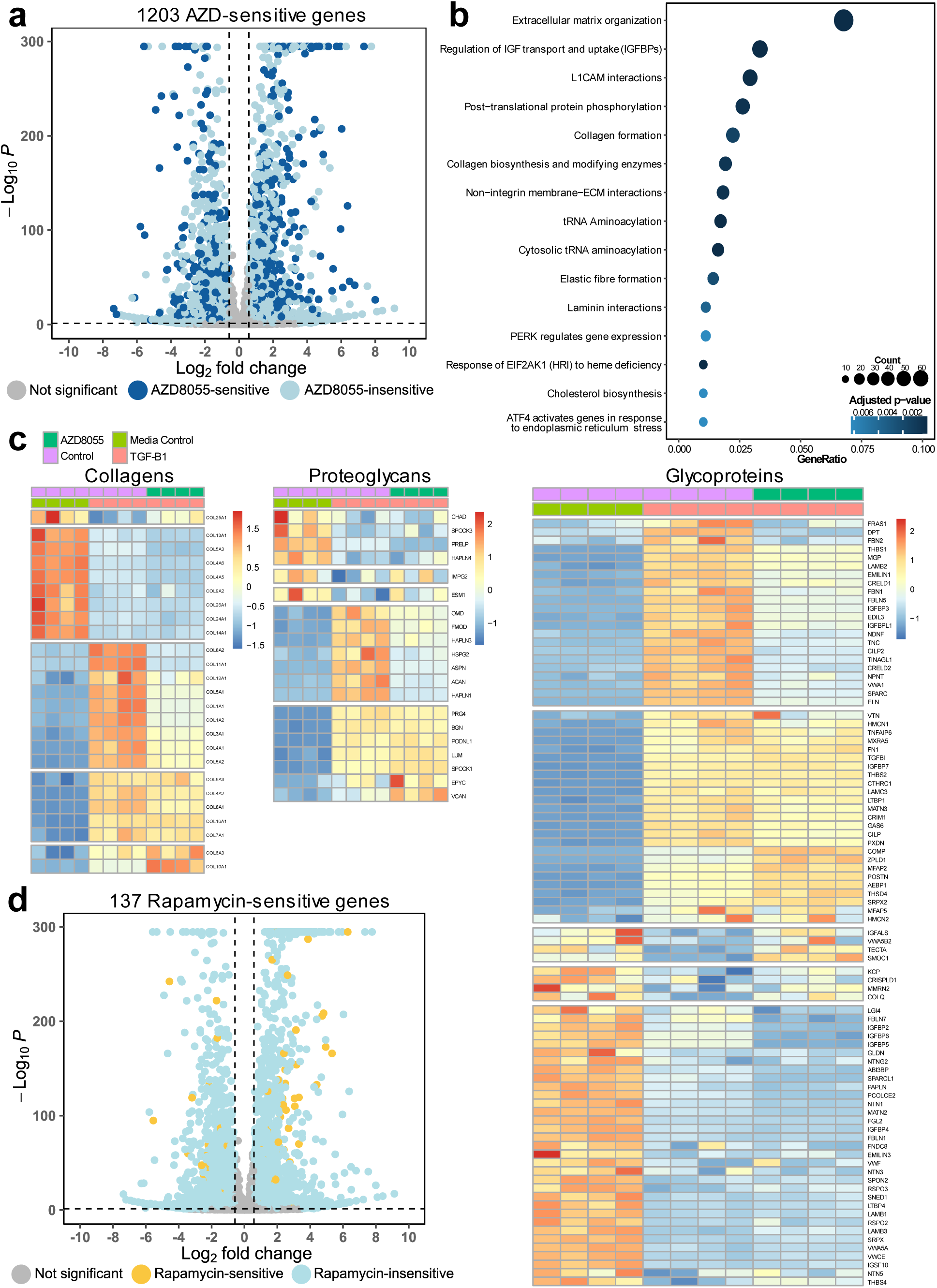
ATP-competitive mTOR inhibition attenuates the TGF-β1-induced transcriptional response in primary human lung fibroblasts. pHLFs were incubated with vehicle (0.1% DMSO v/v), AZD8055 (1 μM) or rapamycin (100 nM) for 1 hour prior to stimulation with or without TGF-β1 (1 ng/ml) for 24 hours and RNA-Seq was carried out. (a) Volcano plot depicting the TGF-β1 response, with 1,203 AZD8055-sensitive genes in dark blue and AZD8055-insensitive genes in light blue (p-adjusted value < 0.05 and fold change > 1.5). (b) Dot-plot showing the top 15 Reactome pathways (p-adjusted value < 0.01) from over-representation analysis of the AZD8055/TGF-β1-dependent genes. Count refers to the number of genes in the data set that are part of that pathway and GeneRatio refers to the ratio of the number of genes within that pathway that are differentially expressed, to the total number of genes in that pathway. (c) Heat-maps representing supervised clustering analysis of collagen, proteoglycan and glycoprotein genes of media control, TGF-β1-stimulated and TGF-β1/AZD8055-treated pHLFs. Colour scale represents normalised counts scaled to Z score. (d) Volcano plot depicting the TGF-β1 response, with 137 rapamycin-sensitive genes in yellow and rapamycin-insensitive genes in light blue (p-adjusted value < 0.05 and fold change > 1.5).

Exploration of the effect of the partial mTORC1 inhibitor, rapamycin, on the TGF-β1-induced ECM in the same dataset (GSE102674) revealed that only 137 TGF-β1-responsive genes were found to be sensitive to rapamycin. This represents just 2.5% of the total TGF-β1-responsive genes (Figure 1d; for a full gene list please see interactive Supplementary Table 3). Pathways analysis of these rapamycin-sensitive DEGs revealed that they are overrepresented in two pathways: ‘Regulation of cholesterol biosynthesis by SREBP (SREBF)’ and ‘Metabolism of steroids’ (Supplementary Figure 1a). Only a very small number of rapamycin-sensitive matrisome genes, with the majority being secreted factors, were identified (Supplementary Figure 1b).

### The TGF-β1-induced pro-fibrotic gene programme is under mTORC1 regulatory control

The large number of AZD8055-sensitive genes identified within the TGF-β1 induced transcriptome, led us to next determine the relative contribution of the two individual mTOR complexes to this response. To this end, we subjected pHLFs to CRISPR-Cas9 gene editing of either *RPTOR* or *RICTOR* to disable mTORC1 and mTORC2 signalling respectively, and performed RNA-Seq in cells exposed to TGF-β1 for 24 hours. Differential gene expression analysis revealed that of the genes regulated at the transcriptional level by TGF-β1, only 13 genes were mTORC2-dependent (for a list of these genes please see interactive Supplementary Table 5) and there were no significant mTORC2-dependent pathways identified through pathways analysis. In contrast, 1,438 of the TGF-β1 responsive genes (Figure 1a) were mTORC1-dependent (Figure 2a; for a full gene list please see interactive Supplementary Table 4). The top over-represented 15 mTORC1-dependent pathways comprise contain those critical to fibrogenesis (Figure 2b), including the top pathway of ‘Extracellular matrix organisation’ with 60 associated genes. The DEGs mapping to these ECM-related pathways are depicted in Figure 2c. When classifying these genes as either matrisome or non-matrisome genes, we found that, of the TGF-β1-dependent matrisome genes, 35% were under mTORC1-dependent transcriptional control (Figure 2d). This includes genes encoding the pro-alpha 1 chain of the two most prevalent collagen types within the ECM, *COL1A1* and *COL3A1*, and other fibrillar collagens (e.g. *COL5A1*, *COL5A2* and *COL11A1*), as well as several collagen cross-linking genes (e.g. *PLOD2*, *LOX*, *LOXL2*). Furthermore, genes which are associated with the mTOR axis were also TGF-β1 and mTORC1 regulated (e.g. *EIF4EBP1, EIF4B, RHEB*), suggesting a potential feedback loop in the regulation of the mTOR signalling hub.

**Figure 2:**
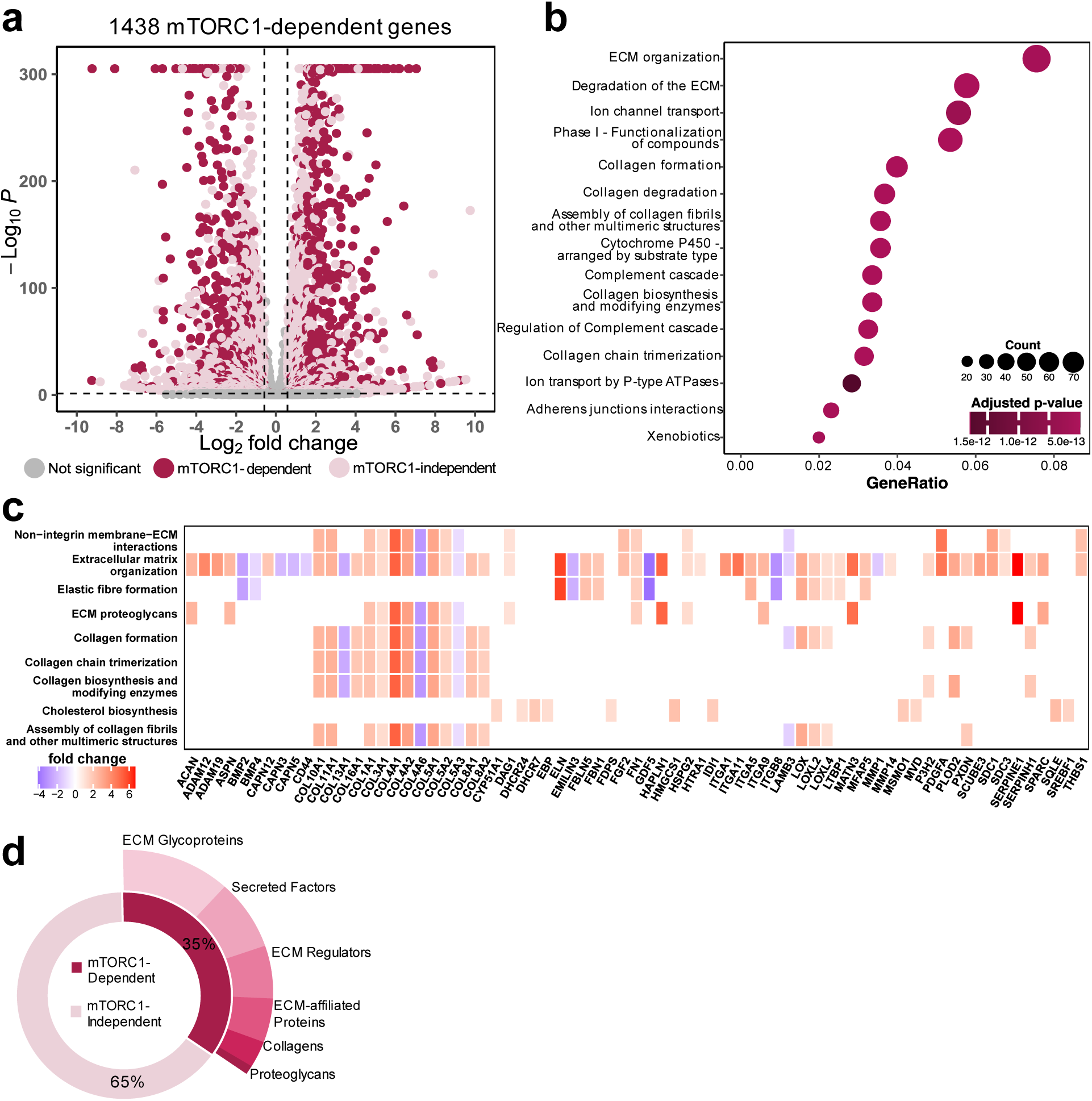
RNA-Seq analysis in combination with CRISPR-Cas9 gene editing reveals that the TGF-β1-induced fibrotic response is under mTORC1 regulatory control. pHLFs were modified using CRISPR-Cas9 gene editing using gRNAs targeting RPTOR or RICTOR and the cells were stimulated with or without TGF-β1 (1 ng/ml) for 24 hours and RNA-Seq was carried out. (a) Volcano plot showing the number of differentially expressed genes following TGF-β1 stimulation (p-adjusted value < 0.05 and absolute fold change > 1.5) with 1,438 mTORC1-dependent genes highlighted in maroon, and mTORC1-independent genes in light pink. (b) Dot-plot showing the top 15 Reactome pathways (p-adjusted value < 0.01) from over-representation analysis on the TGF-β1/mTORC1-dependent genes. Dot size represents the number of enriched genes, and the x-axis represents the fraction of enriched genes over the total number in the Reactome gene set. The colour scale represents the p-adjusted value. (c) Heat-plot showing gene membership of the mTORC1-dependent ECM pathways from the top 15 presented in (b). The colour scale represents the TGF-β1-induced fold change of a given gene within a pathway. (d) Doughnut chart showing the number of TGF-β1-dependent matrisomal genes that are mTORC1-dependent (counter-modulated in RAPTOR cells with p-adjusted value < 0.05 and absolute fold change > 1.5).

### The TGF-β1/mTORC1 axis plays a critical role in defining the transcriptional identity of a high collagen producing *CTHRC1*+ fibroblast subpopulation in IPF

Following the observation that a large number of pro-fibrotic genes in TGF-β1-stimulated cells are regulated by mTORC1, we next explored whether mTOR might play a role in shaping the functional complexity of fibroblast subpopulations in the fibrotic niche. To this end we integrated our RNA-Seq dataset obtained from TGF-β1-stimulated *RPTOR* and *RICTOR* gene-edited pHLFs (Figure 2) with existing, publicly available human IPF scRNA-Seq datasets. As well as the more traditional markers of activated fibroblasts, *COL1A1* and *ACTA2*, (7, 8) novel markers of high collagen expressing fibroblast populations in IPF include *CTHRC1, HAS1* and *PLIN2*. Fibroblasts and myofibroblasts were extracted from the human IPF scRNA-Seq dataset GSE136831, with disease status (IPF = maroon vs control = green) for each cell shown in Figure 3a. Figures 3b-f show the expression of each of the above markers in the reference human scRNA-Seq dataset (GSE136831). This analysis is consistent with previous reports that identified *CTHRC1* as a more specific marker of the fibroblast population that produces the highest levels of ECM proteins in pulmonary fibrosis and other fibrotic diseases (8, 21).

**Figure 3:**
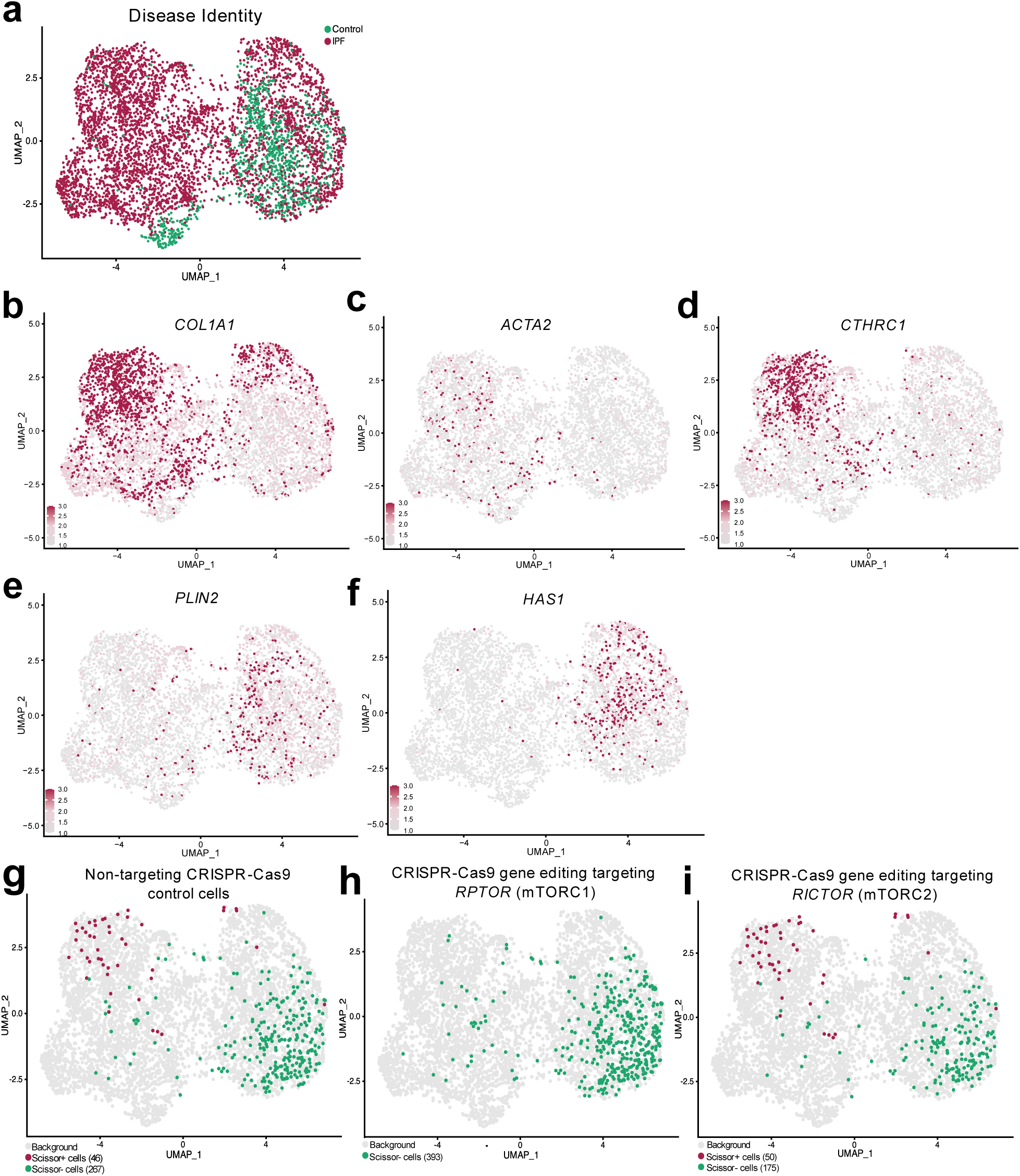
Integration of IPF scRNA-seq and in vitro pHLF RNA-seq data reveals a role for the TGF-β1/mTORC1 axis in defining the transcriptional identity of the CTHRC1+ fibroblast population present in the IPF lung. Scissor was used to map RNA-Seq data (from Figure 2) onto a human IPF scRNA-Seq dataset (GSE136831). (a) UMAP visualisation of the fibroblasts from the scRNA-Seq dataset with the disease status of each cell shown. Control is shown in green and IPF in maroon. (b-f) UMAP visualisation of the expression levels of key activated fibroblast markers from the literature including COL1A1 (b), ACTA2 (c), CTHRC1 (d), PLIN2 (e) and HAS1 (f). (g-i) UMAP visualisation of the Scissor-selected cells for non-targeting CRISPR-Cas9 control cells (g), CRISPR-Cas9 RPTOR gene-edited cells (h), and CRISPR-Cas9 RICTOR gene-edited cells (i), where the maroon and green dots are cells associated with TGF-β1-stimulated (Scissor+) and media control (Scissor-) cells, respectively, while background cells are grey.

We then applied the bioinformatics tool Scissor (22), which is a novel approach that utilises the phenotypes collected from bulk assays to identify the most highly phenotype-associated cell subpopulations from single-cell data. In this setting we first applied this tool to allow the identification of the IPF fibroblast subpopulation with a transcriptome which best matches the TGF-β1-induced transcriptome of pHLFs in non-gene edited control cells (i.e. non-targeting CRISPR-Cas9 control group treated with TGF-β1). We then explored this same question in pHLFs in which either mTORC1 (CRISPR-Cas9 *RPTOR* gene-edited group) or mTORC2 signalling (CRISPR-Cas9 *RICTOR* gene-edited group) is disabled. Scissor analysis revealed that the baseline transcriptome of *in vitro* cultured pHLFs closely matches that of freshly isolated fibroblasts from human lungs characterised by low expression of *CTHRC1* and *COL1A1* (Scissor-cells from the non-targeting CRISPR-Cas9 control group, Figure 3g). In contrast, the transcriptome of TGF-β1-stimulated pHLFs overlaps with cells within a population of fibroblasts unique to IPF, characterised by high expression of *CTHRC1* and *COL1A1* (Scissor+ cells from the non-targeting CRISPR-Cas9 control group, Figure 3g). Strikingly, the transcriptome of TGF-β1-stimulated pHLFs in which mTORC1 signalling has been disabled (Scissor+ cells from the CRISPR-Cas9 *RPTOR* gene-edited group, Figure 3h), no longer maps to the *CTHRC1*+ fibroblast population or any other fibroblast population present in human lungs captured through scRNA-Seq (Scissor+ cells from the CRISPR-Cas9 *RPTOR* gene-edited group, Figure 3h). In contrast, similarly to the non-targeting CRISPR-Cas9 control group (Scissor-cells from the non-targeting CRISPR-Cas9 control group, Figure 3g), the transcriptional profile of baseline *RICTOR* gene-edited pHLFs (Scissor-cells from the CRISPR-Cas9 *RICTOR* gene-edited group, Figure 3i) closely matches that of freshly isolated fibroblasts from human lungs characterised by low expression of *CTHRC1* and *COL1A1*. Moreover, the TGF-β1-stimulated *RICTOR* CRISPR-Cas9 group (Scissor+ cells from the CRISPR-Cas9 *RICTOR* gene-edited group, Figure 3i) in turn maps to the high *CTHRC1+* group, suggesting that mTORC2 is dispensable for the acquisition of the pathological *CTHRC1+* fibroblast phenotype. These findings were corroborated in a second human IPF scRNA-Seq dataset, GSE135893 (Supplementary Figure 2), and are consistent with the notion that mTORC1 signalling is necessary for defining the transcriptional identity of the *CTHRC1*+ pathological fibroblast population in response to TGF-β1-signalling *in vitro* as well as in the fibrotic niche in IPF.

Our next aim was to identify the key pathways and genes which define the transcriptional identity of the *CTHRC1*+ pathological fibroblast population. Differential gene expression analysis was applied to the IPF scRNA-seq data (from GSE136831) comparing the Scissor+ cells from the non-targeting CRISPR-Cas9 control group versus all other fibroblasts in human lung (Figure 3g). We identified 145 genes significantly (adjusted p-value < 0.05) upregulated or downregulated (absolute fold change > 1.5), as shown in Figure 4a (for a full gene list please see the interactive Supplementary Table 6). Pathways analysis of the DEGs revealed that the pathways most associated with the Scissor+ population (non-targeting CRISPR-Cas9 control group) versus all other fibroblasts in human lung were predominantly related to extracellular matrix organisation and collagen synthesis (Figure 4b). We identified 6 genes with the highest positive fold changes that define the transcriptional identity of the *CTHRC1*+ fibroblast population including *CTHRC1* and *COL1A1,* as expected (individual expression levels shown in Figure 4c), as well as *ASPN, POSTN, ADAM12* and *TGFBI* (individual expression levels shown in Figure 4d).

**Figure 4:**
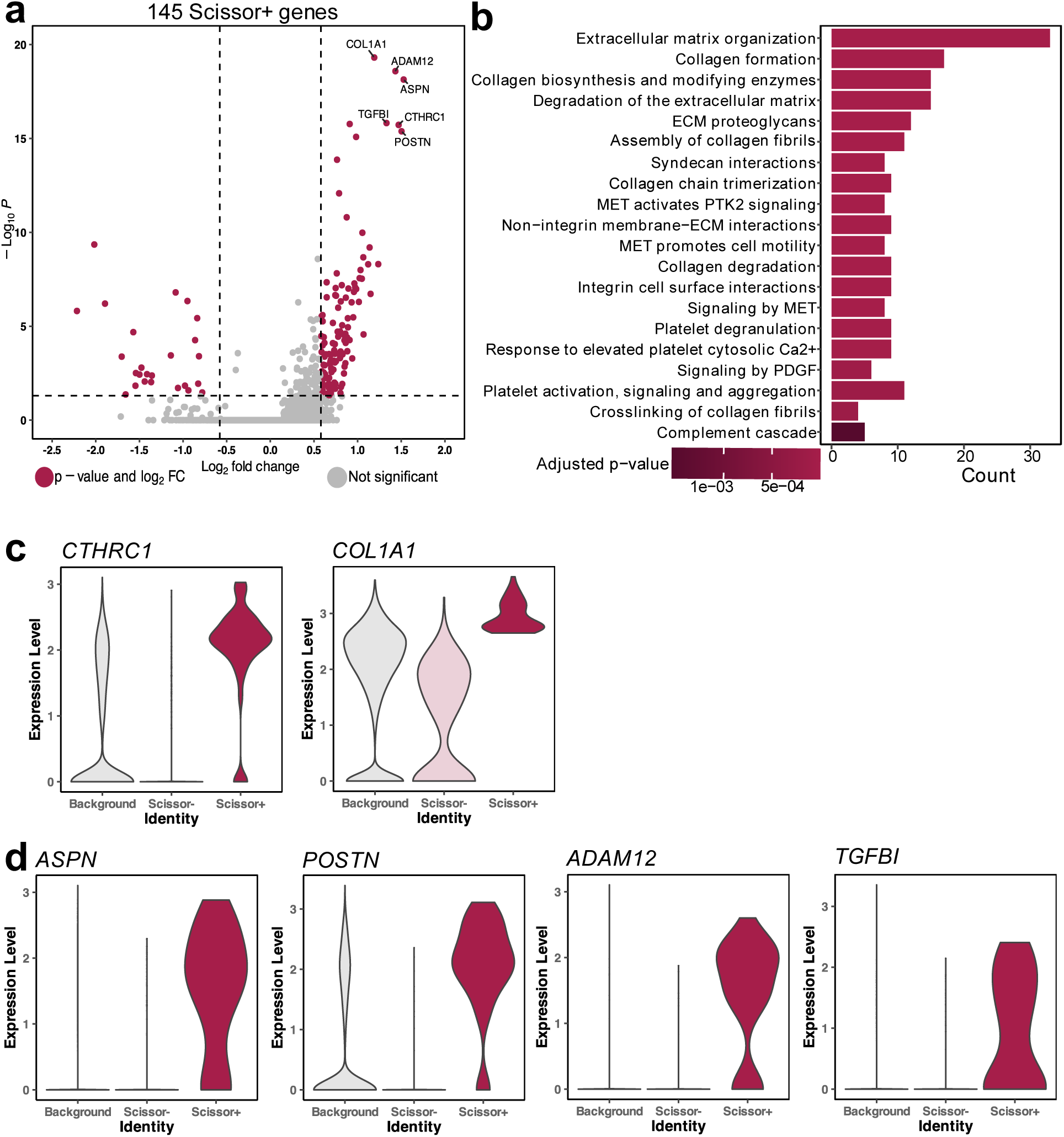
Key pathways and genes defining the transcriptional identity of the CTHRC1+ fibroblast population are dependent on mTORC1 signalling. Scissor was used to map RNA-Seq data (from Figure 2) onto a human IPF single-cell RNA-Seq dataset (GSE136831). (a) Volcano plot highlights the number of differentially expressed genes for the Scissor+ population, when compared to all other fibroblasts. Significant genes are highlighted in maroon (p-adjusted value < 0.05 and absolute fold change > 1.5). (b) Bar-plot showing the top 20 Reactome pathways (p-adjusted value < 0.01) from over-representation analysis of the Scissor+ genes. The x-axis represents the number of genes enriched in that pathway and the colour scale represents the adjusted p-value. (c,d) Violin plots show the expression of the six genes in the Scissor+ population (maroon) with the highest fold changes in comparison to the Scissor-(light pink) and all other fibroblasts in the dataset (grey).

### Selective mTORC1 inhibition prevents the acquisition of a *CTHRC1*+ fibroblast phenotype following TGF-β1 stimulation

We next investigated if there was a direct functional link between mTORC1 signalling and the acquisition of the high collagen expressing *CTHRC1*+ phenotype in response to TGF-β1 stimulation by evaluating the effect of selective mTORC1 inhibition using the novel clinical stage bi-steric mTORC1 inhibitor, RMC-5552, on collagen synthesis and deposition and the expression of the six key marker genes which are associated with the *CTHRC1*+ fibroblast population.

Evaluation of the effect of RMC-5552 on the TGF-β1-induced collagen response revealed that RMC-5552 inhibited TGF-β1-induced procollagen synthesis (IC_50_ = 6.920 x10^-10^ M (95% CI = 3.771 x10^-10^ M to 1.069 x10^-9^ M)) as well as TGF-β1-induced collagen deposition assessed in macromolecular crowding conditions in a concentration-dependent manner (Figure 5a, supplemental figure 3) with no effect of the compound on baseline collagen signals. Consistent with this compound acting via inhibition of mTORC1, western blot analysis demonstrated that RMC-5552 inhibited TGF-β1-induced 4E-BP1 phosphorylation and had no impact on the phosphorylation of the mTORC2 substrate, Akt (Figure 5b). We next examined the role of the TGF-β1/mTORC1 signalling axis on the expression of key *CTHRC1*+ fibroblast population genes by qPCR (Figure 4c, d). All six genes tested were significantly upregulated by TGF-β1 stimulation, and this upregulation was significantly reduced in the presence of an RMC-5552 concentration which had previously shown mTORC1-selective inhibition in primary human lung fibroblasts (Figure 4c). Taken together, these data support a critical role for the TGF-β1/mTORC1 axis in defining the transcriptional identity of the IPF-relevant, highly ECM synthetic, *CTHRC1*+ fibroblast population.

**Figure 5:**
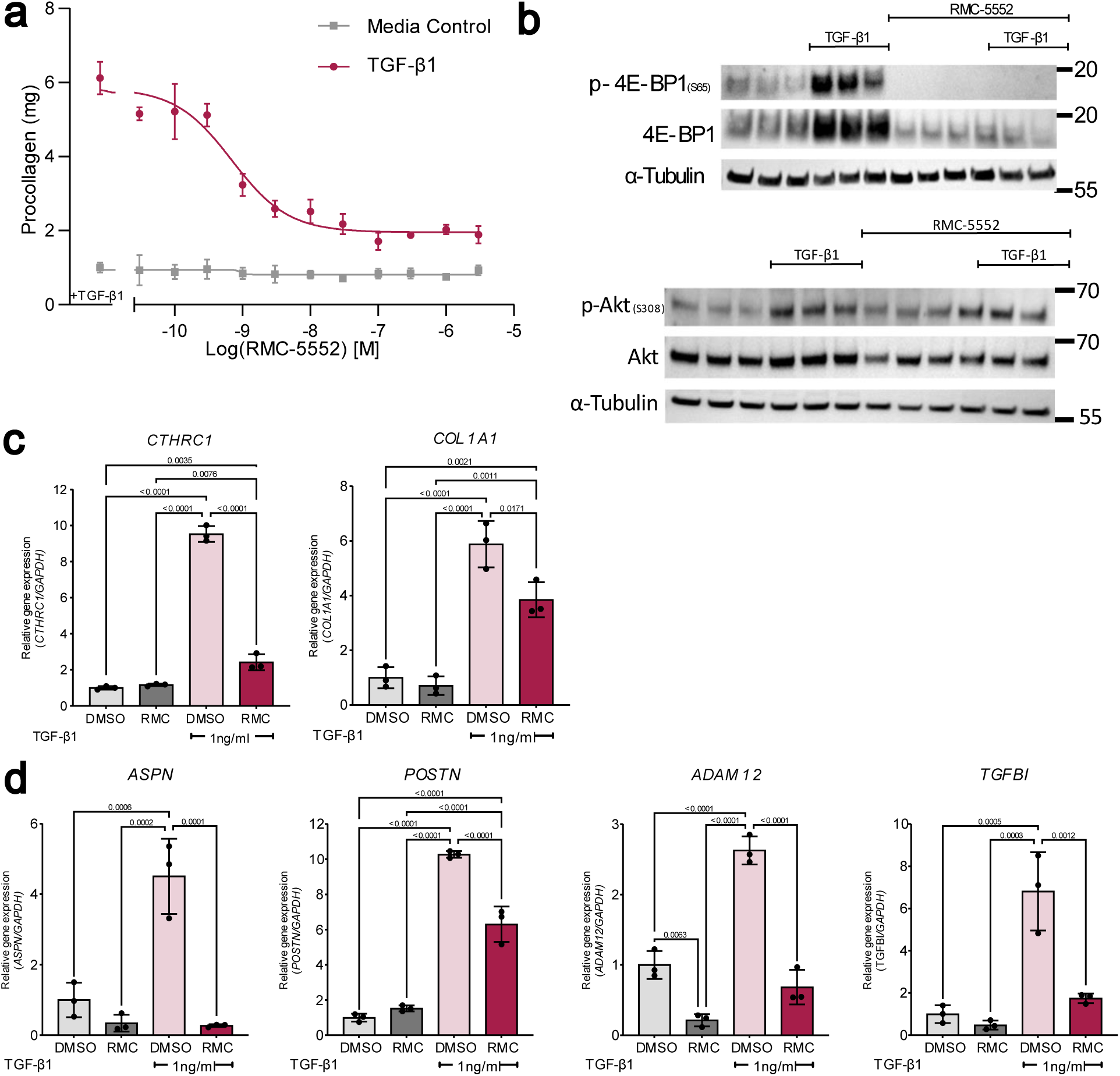
mTORC1-selective inhibitor RMC-5552 inhibits the TGF-β1-induced fibrotic response. pHLFs were exposed to the mTORC1-selective inhibitor RMC-5552 (0.01μM) or a DMSO control and the cells were stimulated with or without TGF-β1 (1 ng/ml) for 24 hours prior to experimental analysis. (a) Hydroxyproline levels were assessed by high-performance liquid chromatography. Each data point shown is mean ± SD (n = 3) and is representative of 3 independent experiments. IC50 values were calculated using a 4-parameter non-linear regression (IC50 = 6.920 x10-10M (95% CI = 3.771 x10-10 M to 1.069 x10-9 M) (b) Phosphorylation of specified proteins is shown by western blot. (c, d) Relative gene expression of the genes (e) CTHRC1, COL1A1 and (f) ASPN, POSTN, ADAM12 and TGFBI are shown by qPCR. Expression is relative to the housekeeping gene GAPDH. Each data point shown is mean ± SD (n = 3) and is representative of 3 independent experiments.

## Discussion

The cardinal pro-fibrotic mediator TGF-β1 has been widely implicated in promoting fibrogenesis in multiple fibrotic conditions. Therapeutic strategies aimed at targeting the TGF-β1 axis in fibrosis, without compromising its critical roles in tissue and immune homeostasis, are therefore actively being pursued. Multiple lines of evidence based on functional *in vitro* and *ex-vivo* studies of human IPF tissue further support a key role for aberrant mTORC1 signalling as a core fibrogenic pathway in IPF and potentially other fibrotic conditions. First, we have previously reported that in addition to the canonical Smad signalling pathway, TGF-β1-induced collagen I production is under critical regulatory control by the mTORC1/4E-BP1 signalling hub and that mTOR signalling acts, at least in part, by influencing *COL1A1* mRNA levels (16). We further identified a key role for mTORC1 in the amplification of the ATF4-dependent de novo serine-glycine pathway to supply glycine during TGF-β1-induced collagen biosynthesis (23). We now report that the role of the mTORC1 axis extends to the regulation of over a third of all TGF-β1 regulated matrisome/matrisome-associated genes and further that the global transcriptome of TGF-β1-stimulated fibroblasts matches that of the *CTHRC1+* pathological fibroblast population recently identified in the fibrotic lung (8). In contrast, the TGF-β1 induced transcriptome of fibroblasts in which mTORC1-signalling is disrupted, does not map to any known fibroblast population. Finally, using the novel clinical stage bi-steric mTORC1 selective inhibitor, RMC-5552, we provide evidence for a direct functional link between mTORC1 and the regulation of key TGF-β1-upregulated genes which define the transcriptional identity of the *CTHRC1+* fibroblast population.

The ECM is composed of a complex array of multidomain macromolecules which are organized in a tissue-specific manner to form a structurally stable network which determines the mechanical properties of tissues. The ECM also acts a reservoir of growth factors and bioactive molecules. Biochemical and biomechanical signals received from the ECM direct cellular function and differentiation and play a decisive role during development as well as the maintenance of tissue homeostasis. The healthy ECM is maintained by the activity of resident fibroblasts and comprises a loose meshwork of collagens, elastin and fibronectin and other core matrisome proteins. During pathological fibrosis, fibroblasts transdifferentiate into activated fibroblasts which synthesize and deposit high levels of ECM molecules and dramatically increase ECM and tissue stiffness by enzymatic covalent crosslinking of collagen and elastin. The profound influence of TGF-β1 on matrisome and matrisome-associated gene expression reported here was not unexpected but the observation that over a third of the TGF-β1-induced ECM transcriptional response is under mTORC1 regulatory control is striking. In contrast mTORC2 is largely redundant for the TGF-β1-induced global transcriptional response in human lung fibroblasts. The mTORC1 regulated ECM transcriptional programme includes numerous core matrisome genes encoding fibrillar and non-fibrillar collagens, elastin, proteoglycans, and other ECM-associated proteins and processing enzymes. Also included are a number of genes encoding glycoproteins which can modulate growth factor signalling, such as fibronectin and members of the fibrillin and latent TGF-β binding protein (LTBP) families which form a microfibrillar network which not only enhances the structural integrity of the ECM, but also influences the binding of growth factors (24–26). We encourage readers to visit our interactive Supplementary Table 1 as a resource to run future queries on these datasets.

In the specific setting of IPF, the histological hallmark of this condition, the fibrotic focus, comprises dense aggregations of activated fibroblasts embedded in a collagen-rich ECM. Recent single cell RNA-Seq studies in IPF have uncovered a large degree of transcriptional heterogeneity among fibroblasts (8, 27), with current evidence indicating the presence of multiple fibroblast populations with unique transcriptomic signatures. These include alveolar fibroblasts, *ACTA2*-positive myofibroblasts, *HAS1*-high fibroblasts, *PLIN2*-positive fibroblasts and *CTHRC1*-expressing fibroblasts (7, 8). Although the relative contribution of these populations to the fibrotic response remains to be fully elucidated, recent evidence indicates that α-SMA is not specific to pathologic ECM-producing fibroblasts (8). In contrast *CTHRC1* was identified as a more specific marker of the small subset of fibroblasts that produce the highest levels of ECM proteins in pulmonary fibrosis and other fibrotic diseases (8). Our analysis using existing IPF datasets is consistent with these observations and shows the highest *COL1A1* and *CTHRC1* expressing fibroblast populations are present in IPF patient lungs, but not in healthy control lungs. The *COL1A1+* and *CTHRC1+* fibroblast population is localised within the collagen rich fibrotic foci in IPF and *cthrc1+* fibroblasts isolated from the lungs of bleomycin-injured mice further exhibit a greater migratory potential, suggesting a role for *CTHRC1+* fibroblasts in the progression of pulmonary fibrosis (8). This fibroblast population has also been identified in other fibrotic conditions, including fibrosis affecting the heart, liver, and skin and constitute a major target for the development of novel anti-fibrotic therapeutic approaches (28–31). While the cellular origin of these subpopulations of fibroblasts in human fibrotic lung disease remains at present elusive, recent lineage-tracing studies in the bleomycin model of lung fibrosis revealed that alveolar fibroblasts represent the main source of multiple fibroblast populations, with the highly ECM-synthetic *CTHRC1*+ fibroblast likely arising from this cell type in response to TGF-β1 signalling within the fibrotic niche (21). Our understanding of the function and role of CTHRC1 during wound repair and fibrosis is at present incomplete. However, CTHRC1 was originally identified as playing an essential role in re-establishing and maintaining tissue homeostasis after wound closure by modulating both the TGF-β and canonical Wnt signalling pathways (reviewed in (15)). This glycoprotein has also been reported to play a protective role based on bleomycin studies in *cthrc1*-knockout mice (32), although the current body of evidence points overwhelmingly to a pathological role for the *CTHRC1+* fibroblast population during fibrogenesis in multiple organs and tissues (31).

The integration of the TGF-β1 regulated transcriptome in human lung fibroblasts using the novel bioinformatic tool Scissor (22), with existing IPF single cell RNA-Seq datasets revealed a key role for mTORC1 signalling in defining the transcriptomic profile of the highly ECM synthetic *CTHRC1+* fibroblast population. We further describe key genes and pathways defining the *CTHRC1+* population with the top pathways impacted in *CTHRC1*+ fibroblasts found to be related to ECM organisation and function. *COL1A1* is unsurprisingly one of the most positively regulated genes within this population but to the best of our knowledge *ASPN, ADAM12* and *TGFBI* have not been previously reported to be highly expressed by the pathologic *CTHRC1*+ fibroblast population. A final key gene, *POSTN,* was recently shown to be highly expressed in *cthrc1*+ mouse fibroblasts (21). *TGFBI* is a known TGF-β-regulated gene and encodes an RGD-containing protein, TGF-β induced protein that plays a role in cell-collagen interactions (33) and has been implicated in the pathogenesis of bleomycin-induced fibrosis (33). *POSTN is* a paralog of *TGFBI and* encodes periostin which plays a multifaceted role in the pathology of fibrotic diseases across various organs fibrosis, including the lung, heart, liver and kidney by modulating cell adhesion, migration, ECM remodelling, inflammation, and fibroblast activation. Moreover, previous work from our laboratory and others has shown that *POSTN* is highly expressed in IPF fibrotic foci and in the bleomycin model of fibrotic lung injury (34, 35). *ASPN* encodes the proteoglycan asporin which has also been reported to be highly upregulated in the lungs of IPF patients (36) and has been found to be localised to *ACTA2*-positive myofibroblasts (37). It influences recycling of the TGF-β receptor and increases its stability at the cell surface. Knockdown of this gene supresses TGF-β1 signalling, which is believed to be due to the loss of this recycling mechanism. It is therefore tempting to speculate that an increase in the expression of *ASPN* in *CTHRC1+* fibroblasts might prolong their responsiveness to TGF-β1 and promote a positive feed-forward loop to drive fibrogenesis to TGF-β1 within the fibrotic niche. *ADAM12* encodes disintegrin and metalloproteinase domain-containing protein 12 and has been implicated in a variety of cellular processes involving cell-cell and cell matrix-interactions. Previous lineage tracing studies in mice have shown that *ADAM12+* cells represent transient precursors to profibrotic cells which produce excessive levels of collagen following tissue injury (38) but the relevance of this observation to human disease remains unresolved.

The data presented here uncover a critical role for the mTORC1 axis in influencing the transcriptional identity of the *CTHRC1+* high collagen expressing pathological fibroblast population induced in response to TGF-β1 signalling within the fibrotic niche. This observation is consistent and extends our previous finding obtained by transcriptome analysis of IPF lung tissue which revealed that both TGF-β1 and the upstream regulator of mTORC1 activation (TSC2/RHEB) formed major signalling clusters associated with collagen gene expression in fibrotic foci captured by laser capture microdissection (34). The importance of mTORC1 signalling in IPF is further supported by genetic association studies. A recent meta-analysis of genome-wide association analysis of five IPF cohorts identified *DEPTOR*, which encodes a major endogenous regulator of mTOR signalling, as a novel IPF susceptibility gene. Importantly, the IPF risk allele is predicted to be associated with reduced expression of DEPTOR and potentially uncontrolled mTOR signalling (39). A second gene in the mTOR pathway, *NPRL3*, which encodes a GATOR1 complex function component and thereby also acts to inhibit mTOR kinase activity has also recently been identified as a potential risk gene in IPF (40).

Our observations have important therapeutic implications in terms of targeting the CTHRC1+ fibroblast population and suppressing excessive ECM deposition in IPF and potentially multiple fibrotic conditions. Therapeutic strategies targeting both the TGF-β1 and mTOR signalling axes have been intensely pursued for over two decades but their success in the clinic has been hampered by the key roles these signalling axes play in immune homeostasis and tumour suppression in the case of TGF-β1, whereas ATP competitive dual mTOR inhibition leads to undesirable effects on glucose metabolism and relief of AKT-dependent feedback inhibition of receptor tyrosine kinase (RTK) expression as a result of concomitant mTORC2 inhibition (19). The recent development of selective bi-steric inhibitors of mTORC1 now offers the potential to overcome some of the major hurdles encountered with these approaches. These mTORC1 selective inhibitors potently inhibit p70S6K and 4E-BP1 phosphorylation, while limiting unwanted effects as a result of concomitant mTORC2 inhibition (19). At the time of writing, the lead compound, RMC-5552, used in this study was being evaluated in early clinical trials in the oncology setting (20). The scientific rationale for evaluating the anti-fibrotic potential of this novel class of selective bi-steric inhibitors of mTORC1 in IPF and potentially other fibrotic conditions is gaining strength.

## Methods

### Primary cell culture

pHLFs were obtained from healthy donors and IPF patients as previously described (16). All cells were cultured at 37°C in 5% CO2 in either Dulbecco’s Modified Eagle Medium (DMEM, Sigma Aldrich) (figures 1-4) or Plasmax (Cancer Tools) (figure 5,6). For cell growth, media was supplemented with 10% fetal bovine serum (FBS) and 1% penicillin streptomycin (both Sigma Aldrich) unless stated otherwise. All experiments were carried out on cells between passages 2 and 8.

**Figure 6:**
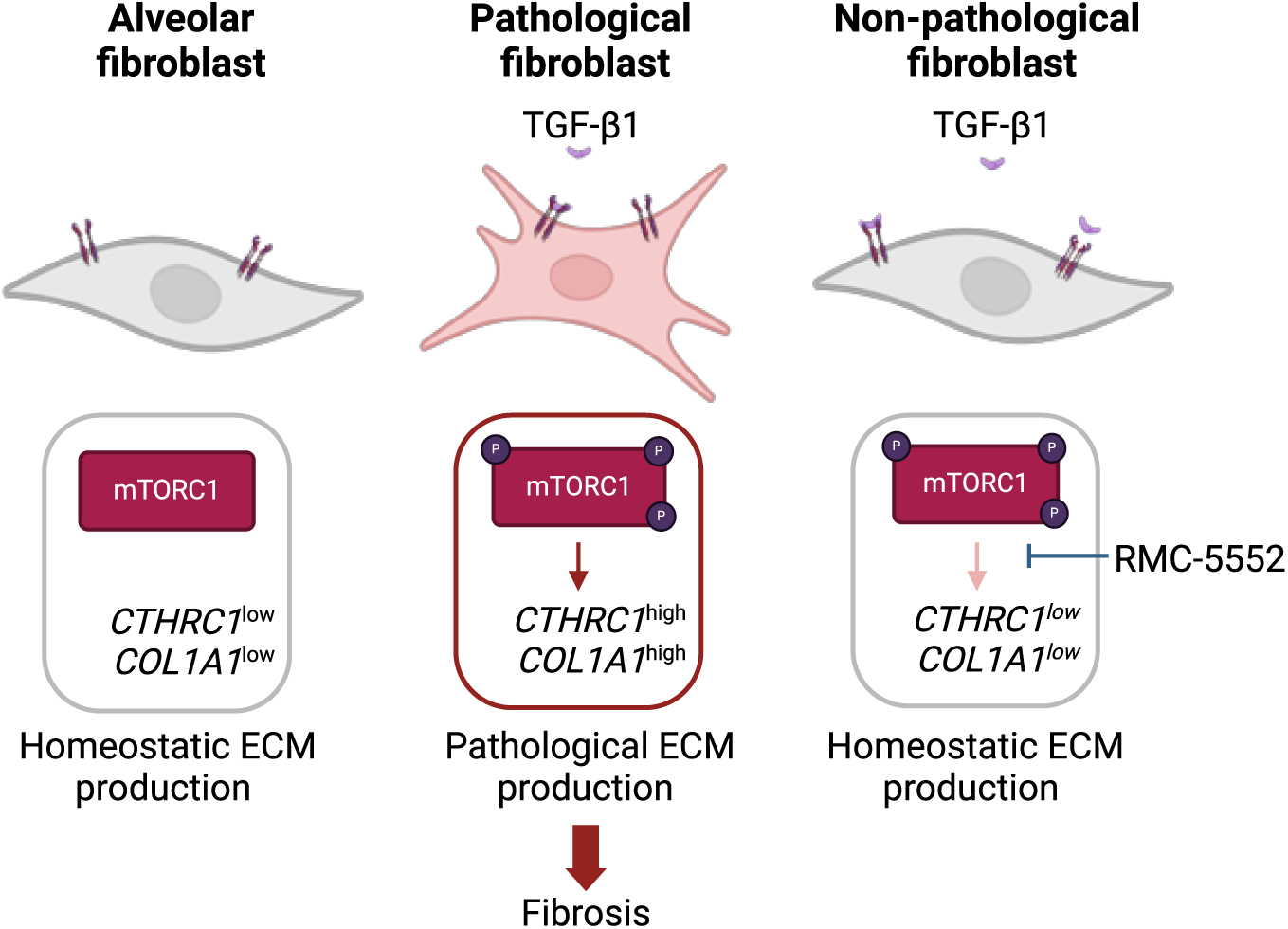
Proposed model of the functional link between mTORC1 and the regulation of key TGF-β1-upregulated genes which define the transcriptional identity of the CTHRC1+ fibroblast population in fibrosis. TGF-β1 activates quiescent fibroblasts and the mTORC1 axis. mTORC1 controls the upregulation of key genes which define the transcriptional identity of the CTHRC1+ fibroblast population, which plays a key role in pathological ECM production and resulting fibrosis. RMC-5552 is an mTORC1-selective inhibitor that can be used to target this pathway.

For TGF-β1 stimulation, cells were exposed to 1 ng/ml TGF-β1 for 24-48 hours depending on the downstream assay. For RMC-5552 exposure, RMC-5552 (HY-132168, MedChem Express) was resuspended in DMSO [1 mM] and diluted to the required concentrations using physiologically relevant media supplemented with 0.4% FBS (Plasmax, Cancer Tools). Cells were exposed to RMC-5552 or a DMSO control for 48 hours.

### CRISPR-Cas9 gene editing

CRISPR-Cas9 gene editing was carried out in pHLFs according to methods described previously (16) Guide RNA (gRNA) sequences were designed using the Deskgen design platform and are outlined in Table 1. Matched wild-type controls were generated using Alt-R Negative Control crRNA 1 (2 nmol, IDT).

**Table 1.**
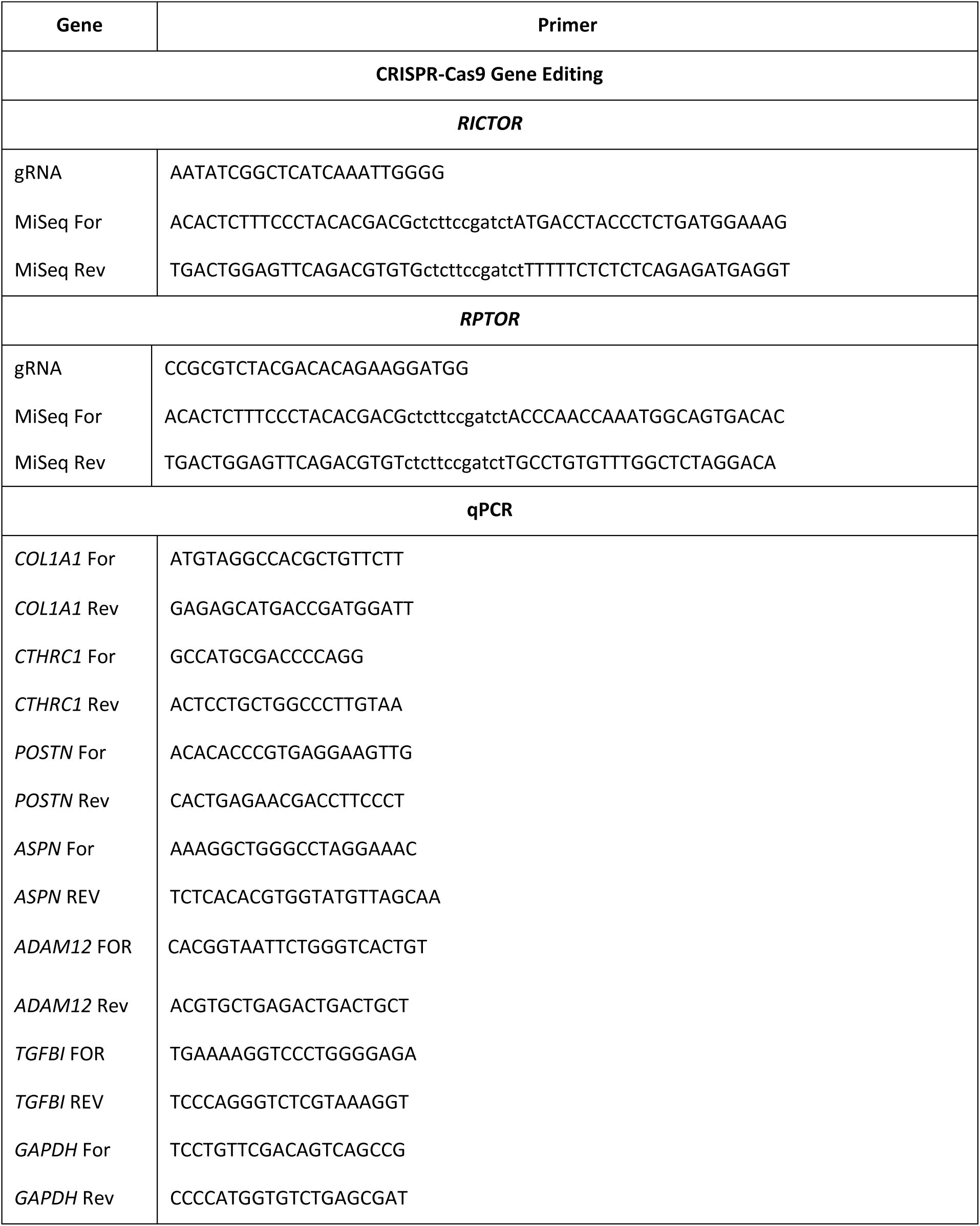
Primers for CRISPR-Cas9 gene editing and qPCR. Rev, reverse. Fwd, forward.

### RNA-Seq

The RNA-Seq datasets described in Figure 1 (GSE102674) were generated as previously described (16). To generate the RNA-Seq datasets described in Figure 2, RNA was extracted using the RNeasy Mini Kit (Qiagen) and 500 ng RNA was used as input for polyadenylate-tailed RNA enrichment and library preparation using the KAPA mRNA HyperPrep Kit (Kapa Biosystems). Assessment of library quality and quantification was carried out using Bioanalyzer High Sensitivity DNA analysis and the Qubit High Sensitivity fluorometry assay according to the manufacturer’s instructions. Paired-end sequencing was performed on the NextSeq sequencing platform (Illumina).

### RNA-Seq analysis

The sequencing data were uploaded to the Galaxy web platform (41), where sequenced reads were quality tested using FastQC and MultiQC (42, 43) aligned to the hg38 human genome using HISAT2 with default parameters (44). Quantification of counts over reference genes was then performed using FeatureCounts (45). Differential gene expression analysis was carried out using DESeq2 (46) within R/RStudio (47, 48) and differentially expressed genes were defined as having an adjusted p-value < 0.05 and fold change > 1.5 or < -1.5 when comparing two experimental conditions. The ClusterProfiler package was used within R/RStudio to conduct over-representation pathways analysis using the Reactome database (49, 50) and significant pathways were defined as having an adjusted p-value < 0.05. Data visualisation was carried out using ClusterProfiler, as well as EnhancedVolcano (51) and pheatmap (52).

### Scissor analysis

The processed scRNA-Seq gene expression matrices were extracted from the Gene Expression Omnibus (GSE136831 and GSE135893). For GSE136831, normalised counts for cells labelled as fibroblasts or myofibroblasts by Adams et al. (27) were extracted. For GSE135893, normalised counts for cells labelled as fibroblasts, *HAS1* high fibroblasts, myofibroblasts and *PLIN2+* fibroblasts were extracted. The Seurat preprocessing function within the Single-cell identification of subpopulations with bulk sample phenotype correlation (Scissor) R package was used to preprocess the data and create a cell-cell similarity network (22). The Scissor algorithm was then applied to identify individual cells within the scRNA-Seq dataset with expression profiles that are correlated with the expression profile of media control or TGF-β1-stimulated samples from the bulk RNA-Seq dataset. The alpha parameter of 0.5 was used for which the reliability significance test reported p < 0.001. The FindAllMarkers function in Seurat was used to identify differentially expressed genes for the Scissor+ cells versus all other cells (p-value < 0.05 and fold change > 1.5 or < -1.5) (53). Pathways analysis was carried out using ClusterProfiler as described above (49, 50).

### Real-time quantitative PCR

Total RNA extraction was carried out using the RNeasy Mini Plus kit (QIAGEN) according to the manufacturer’s instructions. cDNA synthesis was carried out with 500 ng total RNA input using the qScript cDNA Supermix kit (Quanta Biosciences) according to the manufacturer’s instructions. qPCR was performed using the Power SYBR Green PCR Master Mix (Thermo Fisher Scientific) with 5 *µ*l of SYBR green PCR mix per 2 *µ*l of cDNA sample. Forward and reverse primers were added at a final concentration of 800 nM and sequences of primers used are outlined in Table 1. qPCR was performed on the QuantStudio 5 platform (Thermo Fisher Scientific) with the following cycling conditions for 40 cycles: 95°C for 10 minutes, 95°C for 15 seconds and 60°C for 1 minute. The Ct values were normalised to the housekeeping gene *GAPDH* (glyceraldehyde 3-phosphate dehydrogenase) by subtracting the average of the Ct values of the housekeeping genes. Relative expression of genes of interest was then calculated using the 2 deltaCt approach.

### Immunoblotting

Total protein extraction was carried out using ice-cold Phosphosafe buffer (Merck) supplemented with protease and phosphatase inhibitors (Roche). Proteins were run with SDS PAGE and transferred to a membrane using a dry transfer for 7 minutes with an IBlot (Invitrogen). Blocking and antibody washing were performed in 0.5% milk in TBS-T. Protein levels were assessed using the antibodies and dilutions highlighted in table 2. Images were taken with an ImageQuant TL (V8.1).

**Table 2.** Antibodies and dilutions used in Immunoblotting and macromolecular crowding assay.

| Protein | Antibody | Dilution |
| --- | --- | --- |
| Phosphorylated 4E-BP1 (Ser65) | 9451 | 1:2000 |
| 4E-BP1 (total) | 9644 | 1:2000 |
| Phosphorylated Akt (S473) | 4060 | 1:2000 |
| Phosphorylated Akt (T308) | 13038 | 1:2000 |
| Akt (total) | 4691 | 1:2000 |
| Alpha-Tubulin | 9099 | 1:2500 |
| Collagen I | C2456 | 1:1000 |
| Alpha Smooth Muscle Actin | M085101 | 1:500 |
| Alexa-488 | A11001 | 1:1000 |
| Alexa-555 | A21428 | 1:1000 |

### Quantification of hydroxyproline from cellular supernatants

Procollagen production of pHLFs via assessment of hydroxyproline levels was measured by high-performance liquid chromatography as previously described (16).

### Macromolecular crowding assay

Extracellular collagen biosynthesis was measured in 96-well format by a high-content imaging-based macromolecular crowding assay as described previously (16).

### Statistical analysis

All graphs were created in GraphPad Prism (version 10) or within R/RStudio. Where specified in relevant figure legends, data were analysed by one-way ANOVA with Dunnett’s multiple comparisons testing or two-way ANOVA with Tukey multiple comparisons testing. Data were considered statistically significant at *p* < 0.05.

### Study approval

Human samples were acquired following research ethics committee approval (10/H0720/12, 10/H0504/9 and 12/EM/0058) and informed consent was obtained. pHLF cell lines were derived through explant culture, and their purity was confirmed by immunohistochemical characterisation.

## Acknowledgements

The authors gratefully acknowledge the members of UCL Genomics for their contribution to the RNA-Seq aspect of the project. We also thank Adams et al. and Habermann et al. for making the processed scRNA-Seq matrices pre-processed data publicly available to all researchers. We would also like to thank Dr Naftali Kaminski’s group at Yale University for advice on the bioinformatic analyses involved. We acknowledge the support of Chiesi Farmaceutici in providing travel costs for J.A.M.W.

## Author contributions statement

J.A.M.W. and R.W. contributed equally as first authors and performed experiments, analysed and interpreted data, generated figures, and drafted the final manuscript. R.C.C. and M.P. contributed equally as senior authors and conceived, designed, and supervised the study and edited the manuscript. R.C.C. provided funding. D.G. performed experiments. N.K. and S.P. provided expert input into data analysis. All authors reviewed and approved the final submitted manuscript.

## Data availability

The RNA-Seq dataset are deposited and will be publicly available at https://www.ncbi.nlm.nih.gov/geo (accession number will be released upon acceptance).

