## Supplemental materials for "Critical role for the TGF-β1/mTORC1 signalling axis in defining the transcriptional identity of *CTHRC1*+ pathologic fibroblasts"

### Supplemental Tables

The supplementary tables can be found in the uploaded files entitled SupplementaryTables.html.

### Supplemental Figures

**a**

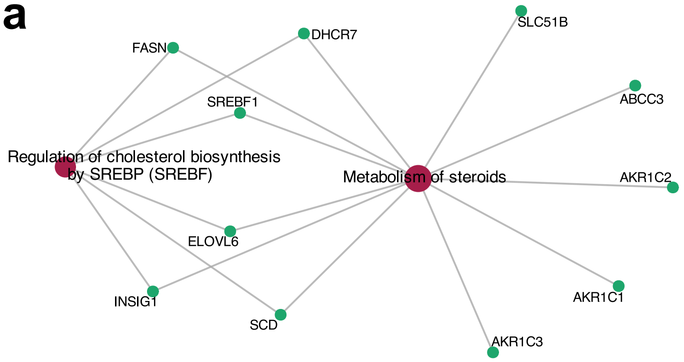


**b**

| TYPE | GENES |
|---|---|
| ECM regulator | *ADAMTS3, ADAMTS6, TGM2* |
| Glycoprotein | *IGFBP5, IGSF10, SMOC1* |
| Proteoglycan | *FMOD* |
| Secreted factor | *ANGPTL4, FGF1, FGF9, FSTL3, GDNF, HGF, IGF1, IL11, NGF, NRG1* |

**Supplementary Figure 1: Rapamycin has limited effects on the TGF-β1-induced transcriptional response in primary human lung fibroblasts.** pHLFs were incubated with rapamycin (100 nM) for 1 hour prior to stimulation with or without TGF-β1 (1 ng/ml) for 24 hours and RNA-Seq was carried out. (a) Gene-connection network plot showing the only two significant Reactome pathways and their member genes (p-adjusted value < 0.01) from over-representation analysis of the rapamycin/TGF-β1-dependent genes. (b) Table showing the limited number of matrisomal genes counter-modulated by rapamycin (p-adjusted value < 0.05 and absolute fold change > 1.5).

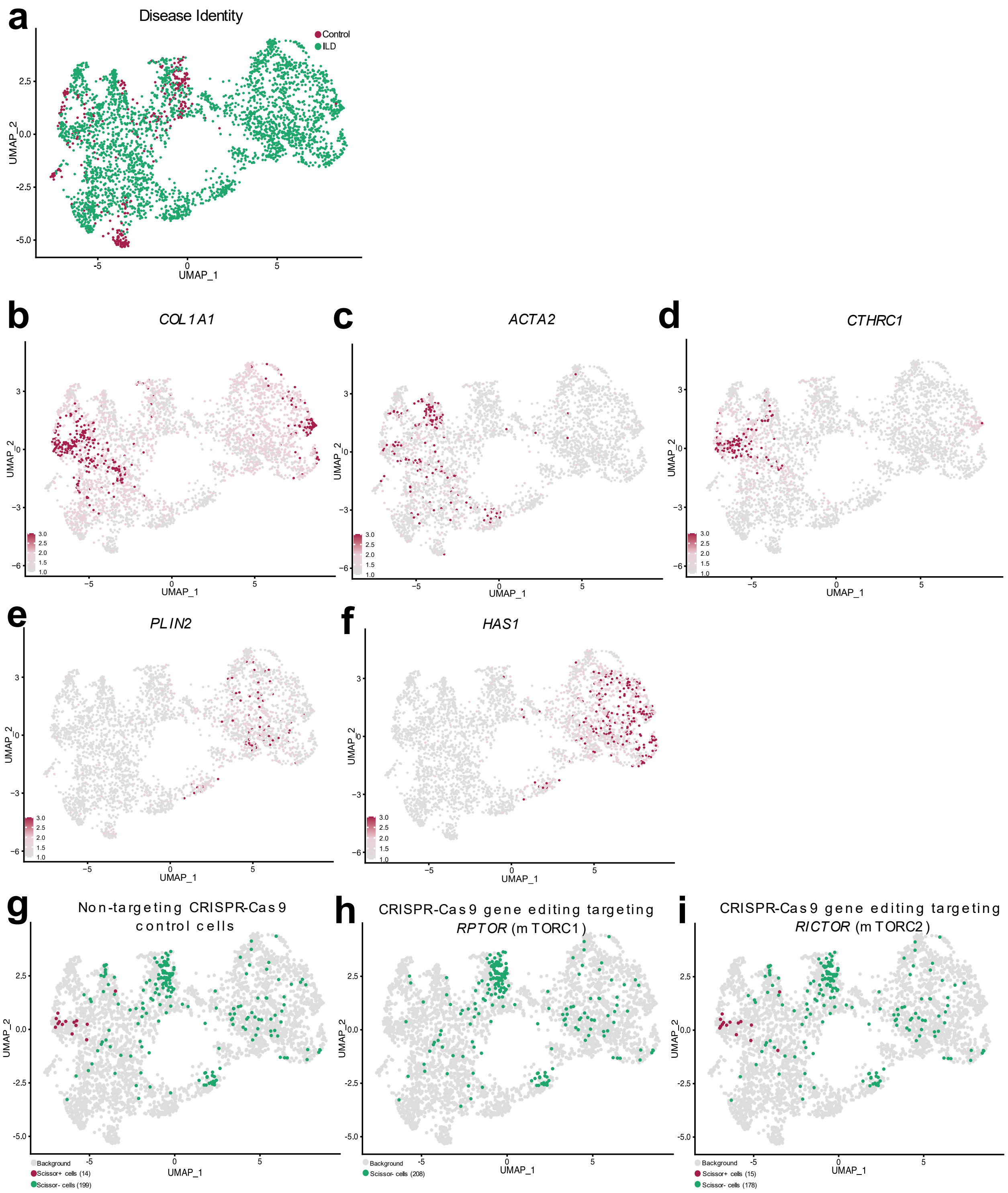


**Supplementary Figure 2: Integration of IPF scRNA-seq and in vitro pHLF RNA-seq data reveals a role for the TGF-β1/mTORC1 axis in promoting the appearance of a CTHRC1+ fibroblast population related to IPF.** Scissor was used to map bulk RNA-Seq data (from Figure 2) onto a human IPF single-cell RNA-Seq dataset (GSE135893). (a) UMAP visualisation of the fibroblasts from the single-cell RNA-Seq dataset with the disease status of each cell shown. (b-f) UMAP visualisation of the expression levels of key activated fibroblast markers from the literature including COL1A1 (b), ACTA2 (c), CTHRC1 (d), PLIN2 (e) and HAS1 (f). UMAP visualisation of the Scissor-selected cells for non-targeting CRISPR-Cas9 control cells (g), CRISPR-Cas9

RPTOR gene-edited cells (h), and CRISPR-Cas9 RICTOR gene-edited cells (i), where the maroon and green dots are cells associated with TGF-β1-stimulated (Scissor+) and media control (Scissor-) cells, respectively, while background cells are grey.

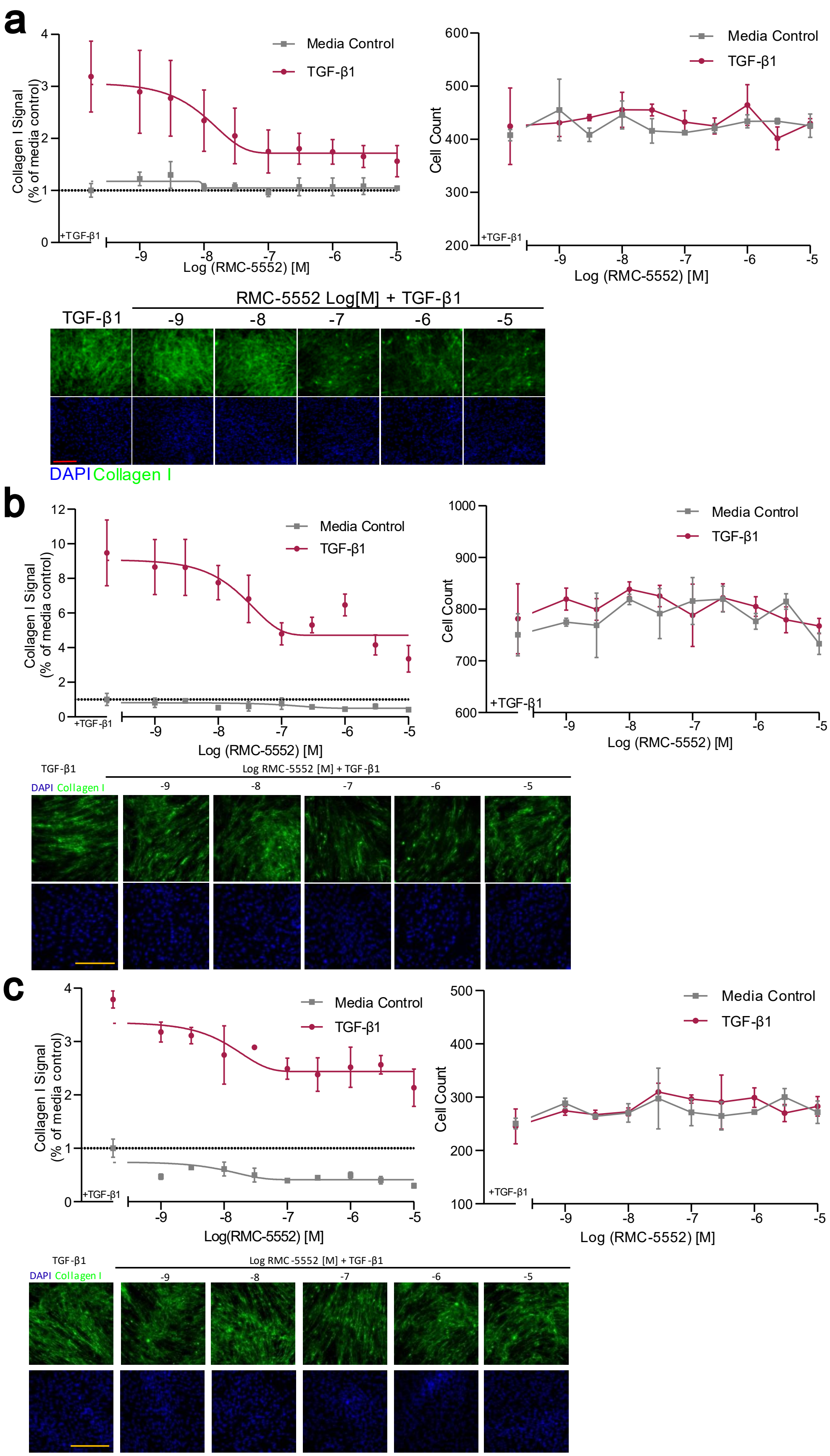
a
Collagen I Signal (% of media control)
Media Control
TGF-β1
+TGF-β1
Log (RMC-5552) [M]
Cell Count
RMC-5552 Log[M] + TGF-β1
TGF-β1
-9
-8
-7
-6
-5
DAPI Collagen I
b
Collagen I Signal (% of media control)
Media Control
TGF-β1
+TGF-β1
Log (RMC-5552) [M]
Cell Count
TGF-β1
Log RMC-5552 [M] + TGF-β1
DAPI Collagen I
c
Collagen I Signal (% of media control)
Media Control
TGF-β1
+TGF-β1
Log(RMC-5552) [M]
Cell Count
Log (RMC-5552) [M]
TGF-β1
Log RMC-5552 [M] + TGF-β1
DAPI Collagen I

**Supplementary Figure 3: mTORC1-specific inhibitor RMC-5552 inhibits the TGF-β1-induced fibrotic response in IPF primary human lung fibroblasts.** pHLFs were exposed to the mTORC1-selective inhibitor RMC-5552 (0.01μM) or a DMSO control and the cells were stimulated with or without TGF-β1 (1 ng/ml) for 24 hours prior to experimental analysis. (a, b, c) Collagen I deposition assessed by macromolecular crowding assay from 1 control (a) and 2 individual IPF patients (b, c) . Data are expressed as the fold change in collagen I signal relative to the media control (4 images per well) and cell counts obtained by staining nuclei with DAPI. Representative images are shown. Each data point shown is mean ± SD (n = 4) and is representative of 3 independent experiments.
