## Supplementary tables for "Critical role for the TGF-β1/mTORC1 signalling axis in defining the transcriptional identity of *CTHRC1*+ pathologic fibroblasts": SupplementaryTables.html


### Supplementary Tables

#### 2024-02-12

#### Summary

This document contains the supplementary data tables for the
transcriptomic analyses described in the paper:

- Pharmacological datasets: RNA-Seq data from pHLFs treated with
  AZD8055 or rapamycin
- CRISPR/Cas9 datasets: RNA-Seq data from CRISPR/Cas9 gene edited
  pHLFs
- Scissor analysis: mapping RNA-Seq data onto a human IPF scRNA-Seq
  dataset (GSE136831)

#### Pharmacological datasets

##### TGF-β1-dependent genes in DMSO control cells

##### AZD8055-sensitive genes

##### Rapamycin-sensitive genes

#### CRISPR/Cas9 datasets

##### mTORC1-dependent genes

##### mTORC2-dependent genes

#### Scissor analysis

##### Genes defining the transcriptional identity of CTHRC1+ fibroblasts
